## Supplementary file1 for "Heparan-6-*O*-endosulfatase 2 promotes invasiveness of head and neck squamous carcinoma cell lines in co-cultures with cancer associated fibroblasts"

**Content**

Figure S1 Sulf2-KO of Cal33 cells using Crispr/Cas9

Figure S2 Targeted Mass spectrometry for the detection of Sulf2

Figure S3 Mono-culture spheroids of HNSCC-WT, Sulf2-KO and HNCAF37 cells

Table S1 Inhibition of Sulf2-mediated desulfation of heparan sulfate by HfFucCS

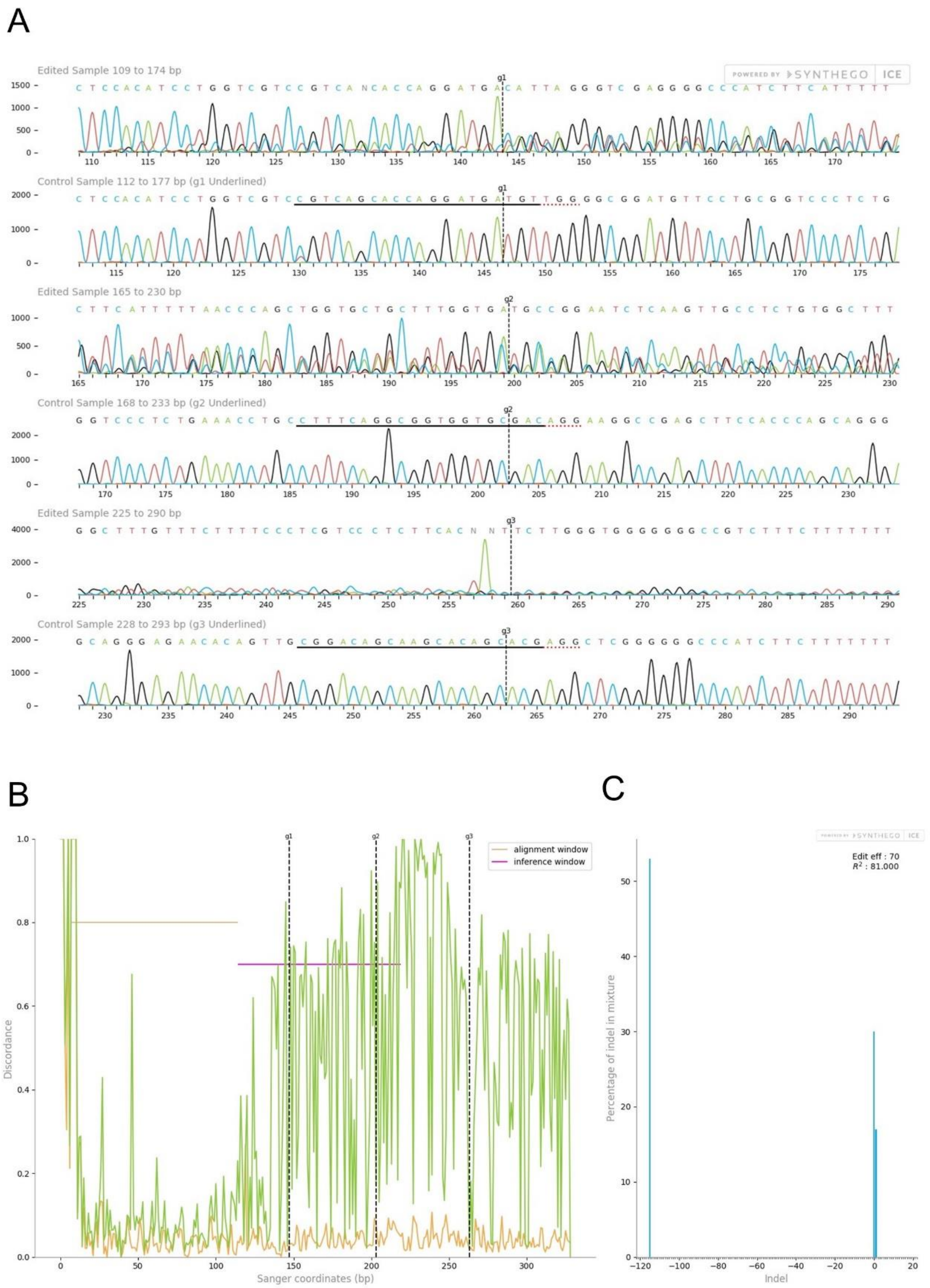
A
Edited Sample 109 to 174 bp
Control Sample 112 to 177 bp (g1 Underlined)
Edited Sample 165 to 230 bp
Control Sample 168 to 233 bp (g2 Underlined)
Edited Sample 225 to 290 bp
Control Sample 228 to 293 bp (g3 Underlined)
POWERED BY SYNTHEGO ICE
B
alignment window
inference window
Discordance
Sanger coordinates (bp)
g1
g2
g3
C
Edit eff : 70
$R^2$ : 81.000
Percentage of indel in mixture
Indel

**Figure S1.** Inference of Crispr Edits (ICE) analysis of Sulf-2 knockout Cal33 cells. (A) Comparison of wild-type (bottom) and knockout (upper) trace files obtained by Sanger sequencing of the edited genomic DNA region. (B) Discordance plot showing the base-by-base level of disagreement between the non-edited wild-type (orange line) and the edited sample (green line) in the inference window around the edited region. (C) Histogram showing the percentual distribution of indel sizes in the edited population. Three sgRNAs are referred to as g1, g2, and g3.

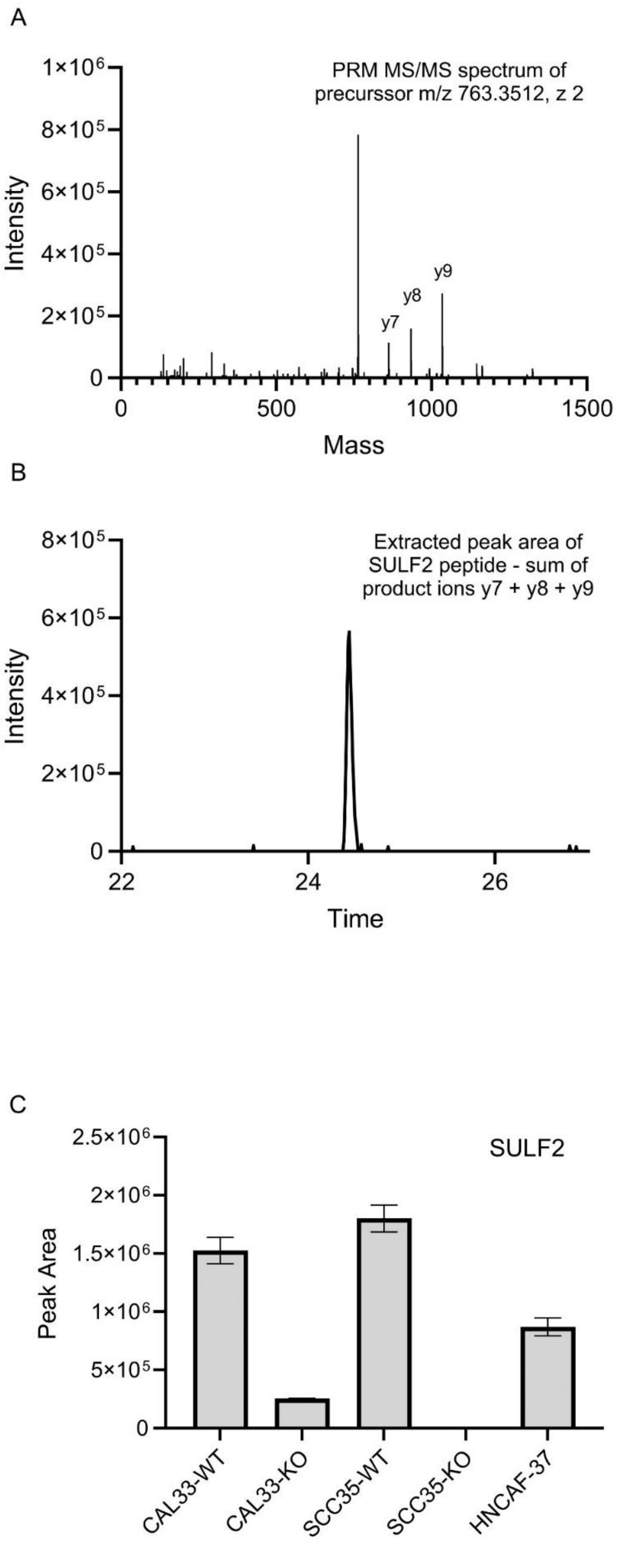
A
PRM MS/MS spectrum of precurssor m/z 763.3512, z 2
Intensity
1×10^6
8×10^5
6×10^5
4×10^5
2×10^5
0
y7
y8
y9
0
500
1000
1500
Mass
B
Extracted peak area of SULF2 peptide - sum of product ions y7 + y8 + y9
Intensity
8×10^5
6×10^5
4×10^5
2×10^5
0
22
24
26
Time
C
SULF2
Peak Area
2.5×10^6
2×10^6
1.5×10^6
1×10^6
5×10^5
0
CAL33-WT
CAL33-KO
SCC35-WT
SCC35-KO
HNCAF-37

**Figure S2.** LC-MS/MS-PRM quantification of Sulf-2 in the secretomes of wild-type and Crispr/Cas knockout cells. (A) Representative PRM spectrum of SCC35-WT sample; the matched Sulf-2 product ions y7 (862.41), y8 (933.44), and y9 (1034.49) are indicated. (B) Extracted peak area of the Sulf-2 peptide using the sum of y7, y8 and y9 product ions at RT 24.44 min. (C) Relative concentration of Sulf-2 in SCC35, Cal33 and HNCAF37 media samples. WT, wild type cells, KO, Sulf-2 KO cells.

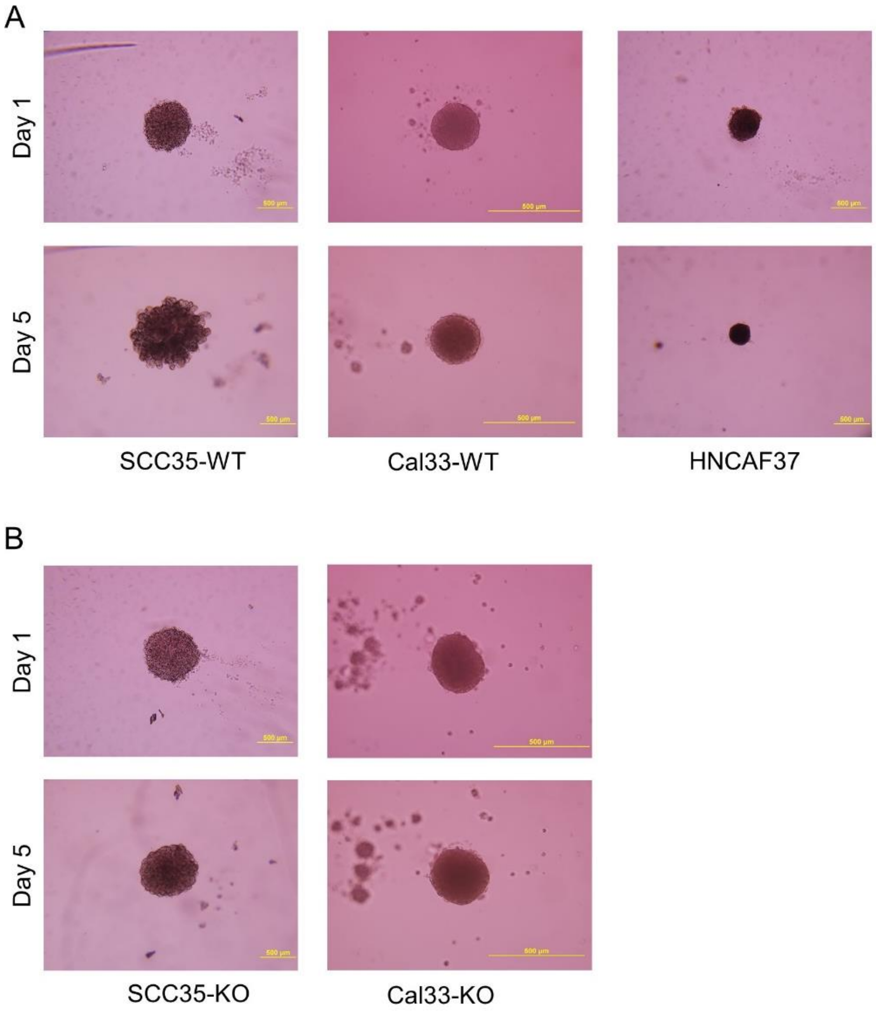


**Figure S3.** Mono-culture spheroids in Matrigel. (A) Representative images of spheroids of HNSCC cells SCC35 and Cal33 and HNCAF37 at day 1 and day 5. (B) Representative images of spheroids formed by SCC35-KO and Cal33-KO at day 1 and day 5. Scale bar 200 µm.

**Table S1.** Inhibition of Sulf-2-mediated desulfation of heparan sulfate by HfFucCS

| Disaccharide | Molar percentage (%) | | |
|---|---|---|---|
| | No treatment | Sulf2 | Sulf2+HfFucCS |
| ΔUA2S-GlcNS6S | 19.4 | 6.0 | 15.4 |
| ΔUA2S-GlcNS | 4.2 | 16.1 | 3.7 |
| ΔUA-GlcNS6S | 16.0 | 11.7 | 12.5 |
| ΔUA-GlcNS | 14.1 | 18.9 | 11.2 |
| ΔUA2S-GlcNAc6S | 1.9 | 0.9 | 1.5 |
| ΔUA2S-GlcNAc | 1.1 | 1.7 | 0.9 |
| ΔUA-GlcNAc6S | 7.3 | 7.7 | 5.7 |
| ΔUA-GlcNAc | 36.1 | 36.9 | 49.0 |

HS from porcine mucosa (25 μg) was desulfated with Sulf-2 (1 μg) with or without HfFucCS (5 μg) for 8 hrs as described in the Methods. Non-treated HS served as a control. The table shows the distribution of individual HS disaccharides as a molar percentage of the sum of all disaccharide forms. Trisulfated disaccharide ΔUA2S-GlcNS6S represents a major Sulf-2 substrate; its desulfation yields ΔUA2S-GlcNS product. Abbreviations: UA, uronic acid; GlcN, glucosamine; GlcNAc, N-acetylglucosamine; S, sulfate.
